# Optimisation of *Ramonda serbica* LEA protein production in *Escherichia coli* and its secondary structure analysis

**DOI:** 10.1101/2025.02.19.639015

**Authors:** Pantelić Ana, Ilina Tatiana, Milić Dejana, Gradišar Helena, Radosavljević Jelena, Marija Vidović

## Abstract

Desiccation, an extreme form of dehydration, reduces the cellular water content to below 5 % and poses a major challenge for most plants. *Ramonda serbica*, a tertiary relict and homoiochlorophyllous resurrection plant, is an exceptional model for investigating vegetative desiccation tolerance. Late Embryogenesis Abundant (LEA) proteins are strongly involved in this adaptive trait, but their exact molecular function is still unclear. In this study, we report the first successful recombinant production of the desiccation-induced LEA protein, RsLEAP30, from a dicotyledonous resurrection plant species using an *Escherichia coli* expression system. By employing immobilised metal affinity and size-exclusion chromatography, we achieved to purify RsLEAP30 to purity over 95 %, providing a robust and scalable method for producing other LEA proteins. Structural characterisation by circular dichroism spectroscopy, combined with *in silico* modelling, revealed that RsLEAP30 is predominantly disordered but contains α-helical regions. We suggest that this structural duality underpins the protective role of RsLEAP30 in chloroplasts, likely via interactions with thylakoids and desiccation-sensitive proteins within photosynthesis-associated proteins. This function may be crucial for the rapid recovery of photosynthetic components upon rehydration. Our study provides new insights into the structure–function relationship of LEA proteins in resurrection plants and establishes a foundation for future investigations. Understanding the protective mechanisms of RsLEAP30 will pave the way for bioengineering strategies aimed at improving the drought tolerance of crops.

## 1. Introduction

Cellular water loss leads to protein denaturation, aggregation, and degradation. It disrupts membrane lipid fluidity, impairs the membrane integrity and function and reduces the cell volume (Oliver et al. 2020). Coping with dehydration is a main challenge for all terrestrial organisms, particularly sessile organisms like plants.

In the course of their evolution and adaptation to terrestrial environments, plants have developed multilevel mechanisms to mitigate the effects of water loss. These strategies include intricate signalling pathways that regulate processes such as stomatal opening and the formation of waxy and suberized cuticles on the surface of vegetative organs (Bowles 2021), allowing most plants to tolerate up to 60 % water loss (Farrant et al. 2007). However, a specialised group of vascular plants, comprising approximately 300 flowering species and known as resurrection plants, has evolved the remarkable ability to survive extreme dehydration or desiccation. When the relative water content drops below 5 %, they enter a state of anhydrobiosis, while they rapidly restore full physiological function after rehydration (Farrant and Hilhorst 2021; Oliver 2020). Among them are three ancient homoiochlorophyllous resurrection plant species from the Gesneriaceae family: *Haberlea rhodopensis, Ramonda nathaliae* and *R. serbica*, which are endemic to the Balkan Peninsula (Rakić et al. 2014). Homoiochlorophyllous plants retain their chlorophyll during desiccation and can quickly resume photosynthesis after rehydration, whereas poikilochlorophyllous plants lose most or all of their chlorophyll during desiccation and have to resynthesise it after rehydration (Georgieva et al. 2007).

Resurrection plants have evolved the unique vegetative mechanisms of desiccation tolerance, which are considered critical steps in the evolution of primitive land plants (Toldi et al. 2009). A key component of their resilience is the upregulation of genes encoding protective proteins, notably heat shock proteins (HSPs) and Late Embryogenesis Abundant (LEA) proteins (Olvera-Carrillo et al. 2011). LEA proteins were first identified four decades ago in cotton seeds (*Gossypium hirsutum*), where they were found to be involved in the late stages of seed maturation (Dure et al. 1981). Subsequent research has shown that they play a broader role in protecting vegetative tissue under environmental stress conditions such as drought, salinity and low temperatures (Battaglia et al. 2008; Hundertmark et al. 2008). LEA proteins are not exclusive to plants. They have also been found in organisms that are tolerant to desiccation, including bacteria and certain invertebrates, underscoring their evolutionary importance for adaptation to stress (Close et al. 1993).

LEA proteins are structurally heterogeneous, predominantly hydrophilic proteins and are considered intrinsically disordered proteins (IDPs) (Amara 2014). IDPs are defined by their lack of stable secondary and tertiary structures under physiological conditions, containing intrinsically disordered regions (IDRs) that enable functional flexibility (Uversky 2019). In their fully hydrated state, LEA proteins primarily adopt random conformations, but during desiccation they undergo to more compact α-helical structures (Hundertmark et al. 2008; Hincha and Thalhammer 2012).

These proteins play a crucial role in stabilising cellular structures, minimising damage during desiccation, and facilitating a rapid return to homeostasis upon rehydration (Olvera-Carrillo et al. 2011; Gechev et al. 2021). LEA proteins may act as molecular shields by mimicking water molecules to protect molecular surfaces during desiccation (Chakrabortee et al. 2017). Additionally, LEA proteins may stabilise cell membranes (Bremer et al. 2017), maintain the integrity of glassy sugar matrices (Hoekstra et al. 2001), chelate redox-active metal ions, and attenuate the accumulation of reactive oxygen species (ROS) (Battaglia 2008). LEA proteins are thought to increase the structural integrity and intracellular viscosity of cells during desiccation by forming separate intracellular proteinaceous condensates (Cuevas-Velazquez et al. 2018; Belott et al. 2020). Recently, it has been shown that the dynamic assembly and disassembly of proteinaceous condensates regulates the availability of free water, ensuring a well-buffered cytoplasmic environment (Watson et al. 2023). Despite all these studies, the exact cellular functions of LEA proteins and their specific intracellular partners remain largely unclear (Dirk et al. 2020).

In our previous studies, we identified 359 LEA proteins from *R. serbica* and classified them into seven classes based on their physicochemical properties (Pantelić et al. 2022). Among them, RsLEA30, a 254-amino acid member of the LEA4 protein group, exhibited a twofold increase in expression and accumulation in desiccated *R. serbica* leaves compared to fully hydrated ones (Vidović et al. 2022). Secondary structure predictions using several algorithms (JPred, Sopma, FELLS, PsiPred, Phyre2) indicate that over 95 % of the amino acid sequence of RsLEA30 has a strong propensity to form α-helices (Pantelić et al. 2022). However, disorder prediction tools showed that about 85 % of the sequence is intrinsically disordered, indicating its structural plasticity under different conditions.

Current efforts to improve drought tolerance in crops focus on biotechnological and synthetic biology strategies that incorporate drought-resistant traits. This emphasises the importance of resurrection plants as model systems for understanding drought tolerance (Farrant and Hilhorst 2021). To fully understand the molecular mechanisms of desiccation tolerance, it is important to understand the physiological function of LEA proteins, which play a key role in this phenomenon.

In this study, we aimed to characterise the structural properties of desiccation-induced RsLEA30 and thereby contribute to a broader understanding of intrinsically disordered stress proteins. We have developed a protocol for the production of pure recombinant RsLEA30 in *Escherichia coli* and investigated its secondary structure *in vitro*.

## 2. Material and methods

### 2.1 Vector, construct and host strain

The expression plasmid pET24a(+) containing the His-tagged GBP-RsLEAP30 construct was sourced from Synbio Technologies (Monmouth Junction, USA), with kanamycin resistance. The construct featured a (His)8 tag followed by a fusion partner, chain A of the immunoglobulin G-binding protein (GBP) from *Streptococcus* sp., followed by an asparagine polylinker (Supplementary Figure S1). The C-terminus of the construct contained a Tobacco Etch Virus (TEV) protease recognition site and the RsLEAP30 gene. The complete construct consisted of (His)_8_-tag-GBP–– polylinker–TEV protease site–RsLEAP30, with NdeI and XhoI restriction sites.

*E. coli* strains DH5α and BL21(DE3) were used for plasmid amplification and protein expression, respectively. After amplification of plasmid in DH5α, the plasmid was validated by sequencing in the Macrogen Europe sequencing facility (Amsterdam, The Netherlands).

### 2.2 Subcloning of RsLEAP30

The vector containing the RsLEAP30 gene with a His tag at the C-terminus was derived from the GBP-RsLEAP30 construct. Primers were designed for amplification: the forward primer contained a NdeI restriction site (5’-CATATGGCGGCGATGTTCGTTGCAAAAACGACCGTGTTAAATTTGTC-3’), and the reverse primer contained an XhoI restriction site, a stop codon and the His_6_ tag, and a (5’-GCAGAGTCTAAACGGCATCACCATCATCACCACTGAATTCTCGAG-3’). The RsLEAP30 gene was amplified by PCR from the ordered plasmid with the mentioned primers. The PCR product was purified using the GeneJET PCR Purification Kit (K0701, Thermo Fisher Scientific) and inserted into a TOPO vector carrying ampicillin and kanamycin resistance markers (11543147, Invitrogen, Waltham, Massachusetts, USA) following the manufacturer’s instructions.

Calcium-competent *E. coli* DH5α cells were transformed by heat shock and plated on LB agar supplemented with kanamycin (50 μg mL^‒1^), ampicillin (100 μg mL^‒1^), IPTG (1 mM), and xGal (20 μg mL^‒1^). Positive, white, colonies were identified by colony PCR and used for plasmid isolation. To ensure proper insertion, empty pET24a(+) and TOPO vectors containing the insert were digested with XhoI, NdeI and HindIII. An additional HindIII digestion confirmed that the insert was cloned. After restriction digestion, the mixtures were purified using the GeneJET PCR Purification Kit, and ligation was performed using a vector: insert ratio of 1:2.5, polyethylene glycol, and T4 ligase according to the manufacturer’s protocol (EL0011, Thermo Fisher Scientific). After incubation at 23 °C for 1 hour, competent DH5α cells were transformed and plated on LB agar containing kanamycin (50 μg mL^‒1^). The presence of the correct insert was confirmed by sequencing plasmids isolated from individual colonies using the GeneJET Plasmid Miniprep Kit (K0503, Thermo Fisher Scientific) and T7 primers in the Macrogen Europe sequencing facility (Amsterdam, The Netherlands).

### 2.3 Small scale RsLEA protein production

For protein expression, calcium competent *E. coli* BL21(DE3) cells were transformed with the pET24a(+) vector containing the construct. A single bacterial colony containing the transformed plasmid was cultured overnight in 10 mL of Luria-Bertani medium (LB) supplemented with 100 μg mL^-1^ of kanamycin and 0.25 % glycerol at 37°C and shaken at 200 rpm.

This overnight culture was then diluted 1:10 into fresh LB supplemented with the 100 μg mL^-1^ kanamycin and 0.25 % glycerol, and grown at 37 °C until the optical density (OD_600_) reached 0.7. An aliquot of non-induced culture was saved as a control, while gene expression was induced using 1 mM isopropyl β-D-1-thiogalactopyranoside (IPTG). Induced cultures were incubated at 25 °C, 30 °C, and 37 °C and shaken at orbital shaker (200 rpm). The aliquots of induced and non-induced broths were collected at various time points (30 min, 1, 2, 3, 4, and 24 hours).

For the optimisation of the expression, cells were harvested by centrifugation at 4000 *g* for 5 min, resuspended in 180 μL of Laemmli buffer, and incubated overnight at 25 °C. The samples were then centrifuged at 15,000 *g* for 10 min, and the supernatants were analysed by SDS-PAGE on 12 % gels as described by Vidović et al. (2016).

### 2.4 Protein concentration

Protein concentrations of the aliquots were determined using the Bradford assay (1976) with a temperature-controlled spectrophotometer (Infinite M Nano+, TECAN, Männedorf, Switzerland) at 595 nm, with measurements performed in triplicate in 96-well microplates.

### 2.5 SDS-PAGE and Western blotting

Proteins were separated by sodium dodecyl sulphate-polyacrylamide gel electrophoresis (SDS-PAGE) on 12 % gels and stained with Coomassie Brilliant Blue G250. For Western blotting, proteins were transferred to polyvinylidene difluoride (PVDF) membranes soaked in Tris-Glycine buffer (pH 8.3) containing 20 % methanol, using a semi-dry transfer method at 1.5 mA cm^‒2^ for 45 min. The blotting was done using SNAP i.d.® 2.0 Protein Detection System for Western blotting (Millipore, Burlington Massachusetts). After blocking with 0.5 % non-fat dry milk in TBST (Tris-buffered saline with 0.05 % Tween 20), the membranes were incubated with primary mouse monoclonal anti-His antibodies (Takara Bio, 631212) diluted 1: 5,000 in TBST with 0.5 % non-fat dry milk for 1 hour. Following four washes in TBST, membranes were incubated with a secondary anti-mouse IgG antibody conjugated to horseradish peroxidase (HRP; Sigma-Aldrich, A9044) diluted 1: 80,000 for 1 hour. After two additional washes in TBST, proteins with His-tags were detected using Immobilon Western Chemiluminescent HRP Substrate (Millipore, Burlington Massachusetts) and visualised with a ChemiDoc Imaging System (Bio-Rad). Protein band quantification on gel was performed using ImageJ (Schneider et al. 2012).

### 2.6 Testing the solubility of GBP-RsLEAP30 and RsLEAP30 recombinant proteins

To assess the solubility of recombinant GBP-RsLEAP30 and RsLEAP30 proteins, *E. coli* cell pellets obtained after incubation under previously optimised conditions, were resuspended in 100 μL of 50 mM Tris-HCl (pH 8) and lysed by sonication for 5 seconds at 10 microns of probe amplitude (Soniprep 150, MSE Crowley, London, UK) on ice. The sonication was repeated three times. The lysates were centrifuged at 15,000 *g* for 10 min at 4°C, and the pellets and supernatants were separated. Both fractions were resuspended in 180 μL Laemmli buffer, incubated overnight at 25 °C, and centrifuged again at 15,000 *g* for 10 min. Equal volumes of the supernatants were analysed by SDS-PAGE.

### 2.7 Large scale GBP-RsLEAP30 and RsLEAP30 protein production

To scale up protein production, the starter, overnight cultures of *E. coli* BL21(DE3) containing the relevant construct used to inoculate 400 mL of LB medium supplemented with kanamycin (50 μg mL^-1^) in 2-liter flasks and shaken at 200 rpm under previously optimised temperature. At an OD_600_ of 0.7, gene expression was induced using 1 mM IPTG induction under optimal conditions. Cells were harvested by centrifugation at 4,000 *g*, resuspended in lysis buffer (1:7 w/v), consisting of 50 mM Tris-HCl (pH 8) and cOmplet EDTA-free Protease Inhibitor Cocktail (Merck, Darmstadt, Germany) and lysed via sonication (12 cycles of 20 s on ice, followed by 20 s pauses). The cell debris was then removed by centrifugation at 15,000 *g* for 20 min at 4 °C.

### 2.8 Protein purification

The lysate was filtered through 0.45 μm and 0.22 μm filters before being purified loaded onto 5 mL HisTrap HP column filed with nickel-based resin (Cytiva, Marlborough, Massachusetts) using the ÄKTA Go™ protein purification system (Cytiva, Marlborough, Massachusetts). A total of 50 mL of filtered lysate was applied to the column pre-equilibrated with 50 mM Tris-HCl (pH 8.00). The target RsLEA proteins were eluted using a stepwise imidazole gradient (10, 20, 30, 40, 50, 75, 100, 150, 200, 250, and 300 mM). Following an optimised elution profile, 2-mL fractions were collected at a flow rate of 2 mL min^‒1^, with absorbance monitored at 280 nm. Imidazole was removed using Amicon® Ultra-15 Centrifugal Filter Units (10 kDa cut-off, Merck). Eluted fractions were analysed by SDS-PAGE. Chosen fractions were pooled, concentrated, snap-frozen in aliquots, and stored at ‒80 °C until use.

The further purification was done on a HiLoad Superdex 75 prep grade resin packed into a 16/100 (17104402, Cytiva, Marlborough, Massachusetts) column size exclusion chromatography (SEC). The resin was pre-equilibrated in 50 mM Tris-HCl (pH 7.5), and protein separation based on molecular weight (Mw) was achieved using the same buffer at a flow rate of 2 mL min^‒1^. The fractions collected were dialyzed against MilliQ water, analysed by SDS-PAGE, lyophilised, and stored ‒80 °C until use. All chromatographic purifications were repeated in at least five independent replicates, with representative chromatograms shown.

### 2.9 Digestion of GBP-RsLEAP30 by TEV

To obtain native RsLEAP30 from the GBP-RsLEAP30 fusion protein, TEV protease (Z03030, GenScript, Piscataway, New Jersey, USA) was used. Following the manufacturer’s instructions, one unit of TEV protease is expected to cleave over 85 % of 3 μg of protein in 50 mM Tris-HCl (pH 8.0) at 30 °C within 1 hour.

### 2.10 Size Exclusion Chromatography Coupled with Multi-Angle Light Scattering (SEC-MALS)

Mw of purified RsLEAP30 was determined using static light scattering with a Dawn Heleos II MALS recorder (Wyatt Technology Europe), coupled to an HPLC system (Alliance e2695, Milford Massachusetts, USA). Selected fractions obtained from SEC were filtered through 0.1 µm centrifugal filter units (Millipore, Burlington Massachusetts, USA), and 50 µL of each sample was injected onto a Superdex 75 Increase 10/300 SEC column (Cytiva, Marlborough, Massachusetts). The proteins were eluted with 50 mM Tris-HCl (pH 7.5). Mw calculations were performed using ASTRA 6 software (Wyatt Technology Europe, Dernbach, Germany), following the manufacturer’s guidelines.

### 2.11 In-gel digestion and MS analysis

Protein bands from gels (1×1 mm) were cut with a clean scalpel, and transferred to tubes. Gel pieces were destained with 10 % acetic acid and 40 % ethanol and 1:1 50 mM bicarbonate and acetonitrile, reduction with 10 mM DTT, and alkylation with 55 mM iodoacetamide were done. After digestion with Trypsin (100 ng), overnight at 37 °C, proteins were extracted with 0.5 % trifluoroacetic acid and 50 % acetonitrile, and dried. Dried samples were resuspended in 2 % acetonitrile, and 0.1 % formic acid in water and injected in Qexactive coupled to a Vanquish Neo High-performance liquid chromatography (HPLC) system (Thermo Fisher Scientific). The samples were loaded on a PepMapTM 100 C18, length 150 mm, internal diameter 0.1 mm, particle size 5 μm (Thermo Fisher Scientific, cat. number 164199) for 5 min with 0.1 % formic acid in water. Two buffers systems were used to elute the peptides: 0.1 % formic acid in water (buffer A) and 0.1% formic acid in 80% acetonitrile (buffer B). Peptide separation was performed using gradient from 2.0 % to 2.0 % in 1.0 min, step 2: from 2.0 % to 37.5 % in 41.0 min, step 3: from 37.5 % to 100.0 % in 2.0 min, step 4: from 100.0 % to 100.0 % in 10.0 min, step 5: from 100.0 % to 2.0 % in 1.0 min, step 6: from 2.0 % to 2.0 % in 5.0 min, of buffer B. Database Search was performed using X!Tandem (version 201.2.1.4, Craig and Beavis 2004).

### 2.12 CD spectroscopy

Circular dichroism (CD) spectra were recorded using a Chirascan CD spectrometer (Applied Photophysics, Leatherhead, UK), equipped with a Peltier temperature controller. To prepare the samples, the buffer in the purest RsLEAP30 fractions was exchanged for MilliQ water. The samples were then freeze-dried and resuspended in MilliQ water to a concentration of 178 μM, which served as the stock solution. For CD measurements, protein solutions were prepared by diluting the stock solution in 10 mM sodium phosphate buffer (pH 4.0-8.0) to achieve a final concentration of 5 μM. Spectral data were recorded from 240 to 190 nm in 1 mm quartz cuvettes (Hellma, Mühlheim, Germany) at 25°C using a 1 nm step size, 1 nm/s scanning speed, 1 s sampling intervals and 1 nm band width. The final spectra represent the average of three scans. The secondary structure content of RsLEAP30 was estimated using three widely used software tools: K2D3 (Louis-Jeune et al. 2012), BeStSel (Micsonai et al. 2022), and DichroWeb (Miles et al. 2021).

### 2.13 3D modelling of RsLEAP30

The sequence of RsLEAP30 was submitted as input for structural prediction using ColabFold (Mirdita et al. 2021). The protein structure was predicted using AlphaFold2 (Jumper et al. 2021). The resulting 3D structure was visualised using BIOVIA Discovery Studio 2019 molecular visualisation software.

## 3. Results

### 3.1 Expression of GBP-RsLEAP30 on a small-scale

The expression of GBP-RsLEAP30 (theoretical Mw = 37 kDa) in *E. coli* BL21(DE3) was confirmed by Western blot analysis with anti-His tag antibodies. A dominant band corresponding to approximately 46 kDa was detected when protein expression was induced by 1 mM IPTG (Figure 1A). A negative control resulted in a weak band corresponding to GBP-RsLEAP30.

**Figure 1.**
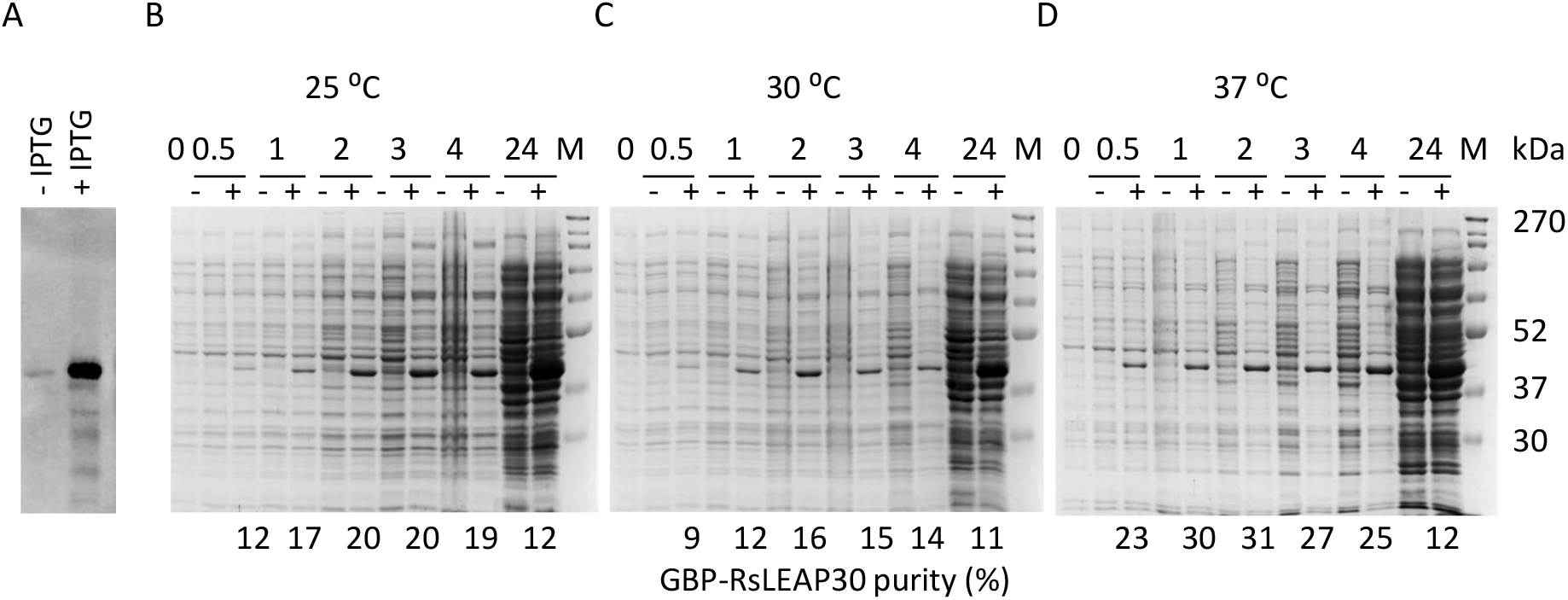
Production of GBP**-**RsLEAP30 in *E. coli* BL21(DE3) in the presence or absence of IPTG inducer. **A)** Western blot of GBP**-**RsLEAP30 with anti-His antibody, after 4 h induction at 37 °C. Optimisation of GBP-RsLEAP30 production over a period of 24 hours at 25 °C **(B)**, 30 °C **(C)**, and 37 °C **(D)**. Equal volumes of total protein extracts were loaded onto gels, which were subsequently stained and scanned according to a standardised procedure. The amounts of GBP-RsLEAP30 and total *E. coli* proteins were quantified using ImageJ. 0, samples before induction; “‒” conditions without IPTG addition; “+” induction with 1 mM IPTG; M, molecular markers (MWP06, NIPPON Genetics, Düren, Germany). The purity of GBP-RsLEAP30 is expressed as a ratio of GBP-RsLEAP30 to total *E. coli* proteins.

To determine the optimal conditions for GBP-RsLEAP30 expression in *E. coli* BL21(DE3), production was analysed at three different temperatures (25 °C, 30 °C and 37 °C) over a 24-hour period. The accumulation of GBP-RsLEAP30 was detectable already within 30 min at all tested temperatures (Figures 1B, 1C, 1D, Supplementary Figure S2). As expected, the highest total yield of GBP-RsLEAP30 was achieved after 24 h, particularly at 25 °C. However, the amount of total *E. coli* proteins was also significantly increased after 24 h at all temperatures. The initial rate GBP-RsLEA30 production was the greatest at 25 °C (Supplementary Figure S2), where after 3 hours of incubation the amount of GBP-RSLEA30 reached 33.1 % of the maximum amount of GBP-RsLEAP30 produced (after 24 h at 25 °C). At 30 °C after 2 hours of incubation, the amount of GBP-RsLEAP30 reached 20.3 %. However, the production rate of GBP-RsLEAP30 at 30 °C decreased over the following 2 hours (Figure 1C, Supplementary Figure S2). Based on the obtained amounts of GBP-RsLEAP30 and the ratio of GBP-RsLEAP30 to total *E. coli* proteins results (Figure 1, Supplementary Figure S2) a 4-hour incubation at 37 °C was selected as the optimal condition for the production of GBP-RsLEAP30 in *E. coli* BL21(DE3).

### 3.3. Solubility test and purification of GBP-RsLEAP30

The protein purification protocol varies depending on whether the protein is soluble or forms inclusion bodies. To assess the solubility of intracellularly synthetized GBP-RsLEAP30, the cell pellets were resuspended in Tris-HCl buffer, lysed and GBP-RsLEAP30 was extracted from the pellet and the supernatant. These fractions were then separated and stained (Figure 2). GBP-RsLEAP30 was detected exclusively in the supernatant (indicated by bands at ∼46 kDa) and not in the pellet, suggesting that GBP-RsLEAP30 is soluble in *E. coli* BL21(DE3) cells.

**Figure 2.**
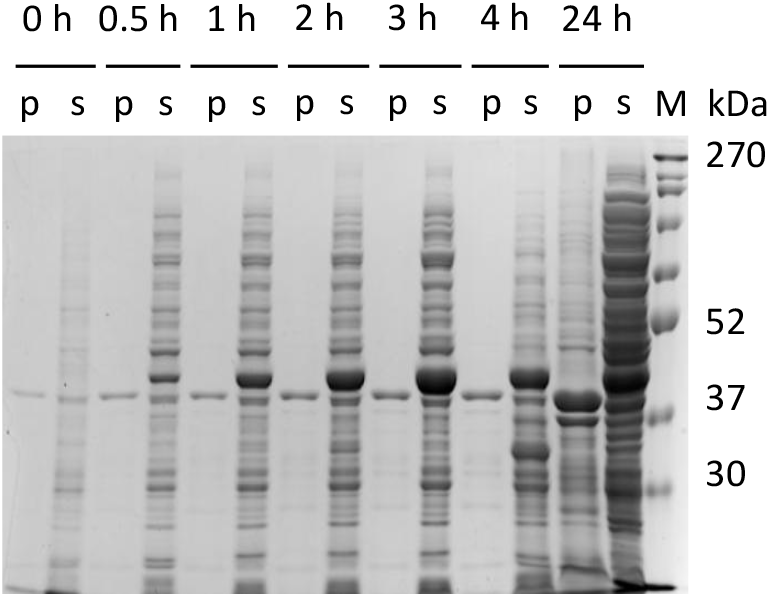
Solubility analysis of GBP-RsLEAP30 during 24 hours of incubation. For details see Materials and Methods, Section 2.5. M, molecular markers (MWP06, NIPPON Genetics, Düren, Germany); p, pellet; s, supernatant.

Given that GBP-RsLEAP30 is His_8_ tagged, the first step of its purification involves immobilised metal affinity chromatography (IMAC). An isocratic elution with 300 mM imidazole was chosen as the most efficient method to elute GBP-RsLEAP30 (Figures 3A, 3B). Remarkably, a significant fraction (70.4 %) of the applied GBP-RsLEAP30 was not retained by the column, as shown by the strong band (∼46 kDa, Figures 3B, 3C) in the flow-through (FT) fraction (Table 1).

**Table 1.** Purification table of GBP-RsLEAP30 from *E. coli* obtained from 1 L culture (from 6 g cell pellet).

| Fraction | Total protein content (mg) | GBP-RsLEAP30 purity (%)<sup>a</sup> | GBP-RsLEAP30 content (mg) | GBP-RsLEAP30 yield (%) |
| --- | --- | --- | --- | --- |
| Lysate | 163.5 | 63.3 | 103.5 | 100.0 |
| Flow through (FT) | 144.7 | 62.5 | 90.4 | 87.4 |
| 300 mM imidazole<sup>b</sup> | 16.8 | 92.7 | 15.4 | 14.8 |

<sup>a</sup> The purity of GBP-RsLEAP30 at each step of purification was estimated by SDS-PAGE.
<sup>b</sup> Polled 300 mM imidazole fraction shown in Figure 3B.

**Figure 3.**
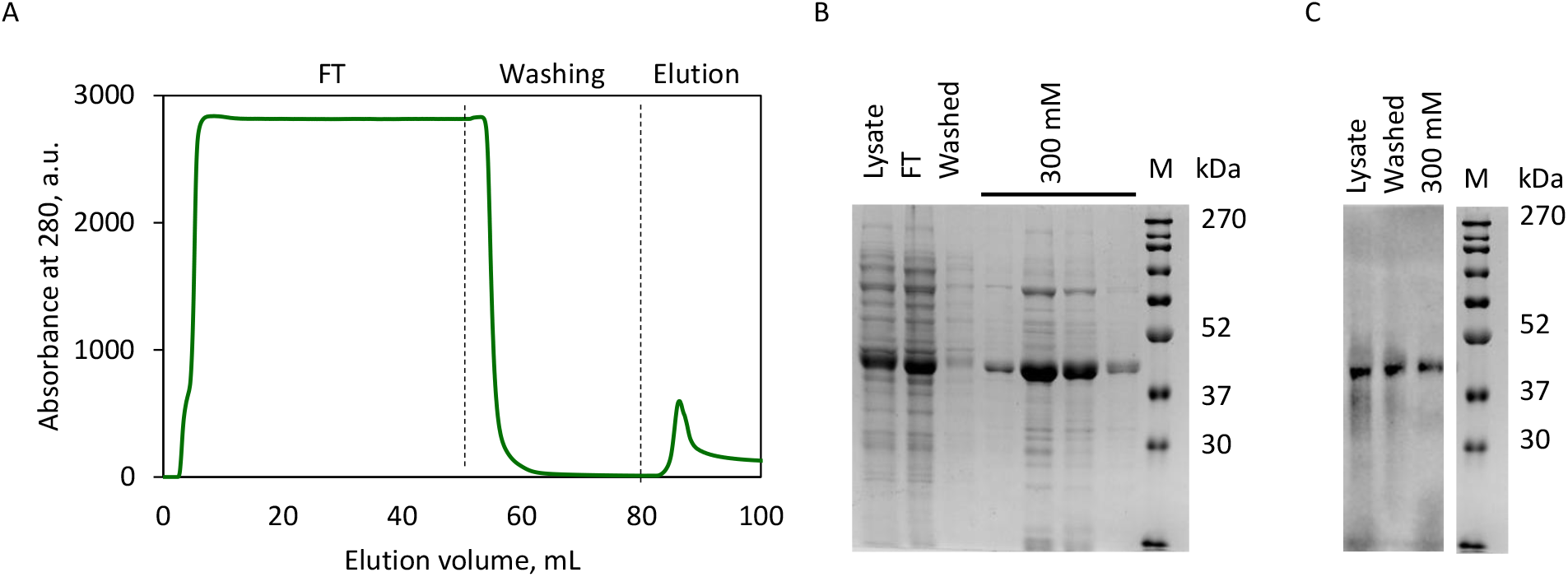
Purification of GBP-RsLEAP30. **A)** Representative IMAC chromatogram of RsLEAP30 elution. The peak containing RsLEAP30 was eluted with 300 mM imidazole. **B)** SDS-PAGE gel of the collected fractions from IMAC. M, molecular markers (MWP06, NIPPON Genetics, Düren, Germany); FT, flow through. **C)** Western blot analysis of the IMAC fractions with anti-His antibody. M, molecular markers (MWP06, NIPPON Genetics, Düren, Germany). The purity of RsLEAP30 is expressed as a percentage of the amount of RsLEAP30 in ratio to the amount of the total *E. coli* proteins.

In total, approximately 15 % of the GBP-RsLEAP30 applied on the column (from 1 L of cell suspension and 6 g of biomass) was recovered with 300 mM imidazole, corresponding to 15.4 mg. The purity of GBP-RsLEAP30 in these fractions was 92.7 % (Table 1).

### 3.4. Digestion of GBP-LEAP30

To remove the fusion partner and His-tag (expected Mw: 9.6 kDa) from GBP-RsLEAP30 and obtain native RsLEAP30 (expected Mw: 27.3 kDa), TEV protease digestion was attempted. Various factors, including pH (7-8), temperature (4-37 °C), incubation time (2-24 h), and the presence of different detergents: 0.05-0.1 % Triton X-100, 0.05-0.1 % Tween 20, 0.05-0,1 % dodecylmaltoside (DDM); 0.1 M urea; reducing agents: 0.6-15 mM cysteine, 0.5-1 mM cystine, 1-20 mM dithiothreitol (DTT), 0.6–20 mM reduced glutathione (GSH), 0.15–0.50 mM oxidized glutathione (GSSG) and their combinations, as well as different molar ratios of enzyme to substrate (1:1, 1:2, 1:5, 1:6, 1:25) and ethylenediaminetetraacetic acid, 0.5-2 mM EDTA, were systematically tested. Despite these extensive optimisations, efficient proteolysis was not achieved, although TEV protease activity was confirmed with other recombinant proteins containing its recognition site (data not shown).

Given the inefficiency of TEV digestion, an alternative strategy was pursued. A new RsLEAP30 construct was designed and subcloned to allow expression of RsLEAP30 containing only a His-tag at the C-terminus, eliminating the need for post-expression cleavage.

### 3.5. Small-scale production of RsLEAP30

The expression of RsLEAP30 (theoretical Mw: 28 kDa) in *E. coli* BL21(DE3) was verified by Western blot with an anti-His antibody (Figure 4A). When analysed by SDS-PAGE, the overexpressed RsLEAP30 routinely displayed three bands (Figures 4B, 4C, 4D, 4E) The higher mobility band (RsLEAP30_H_) corresponded to 31.2 kDa, the intermediate mobility band, RsLEAP30_M_, corresponded to a Mw of 29.5 and the lower mobility band (RsLEAP30_L_ isoform) corresponded to a Mw of 25.6 kDa. These bands were absent in the non-induced control, and were all visualised by anti-His antibody.

**Figure 4.**
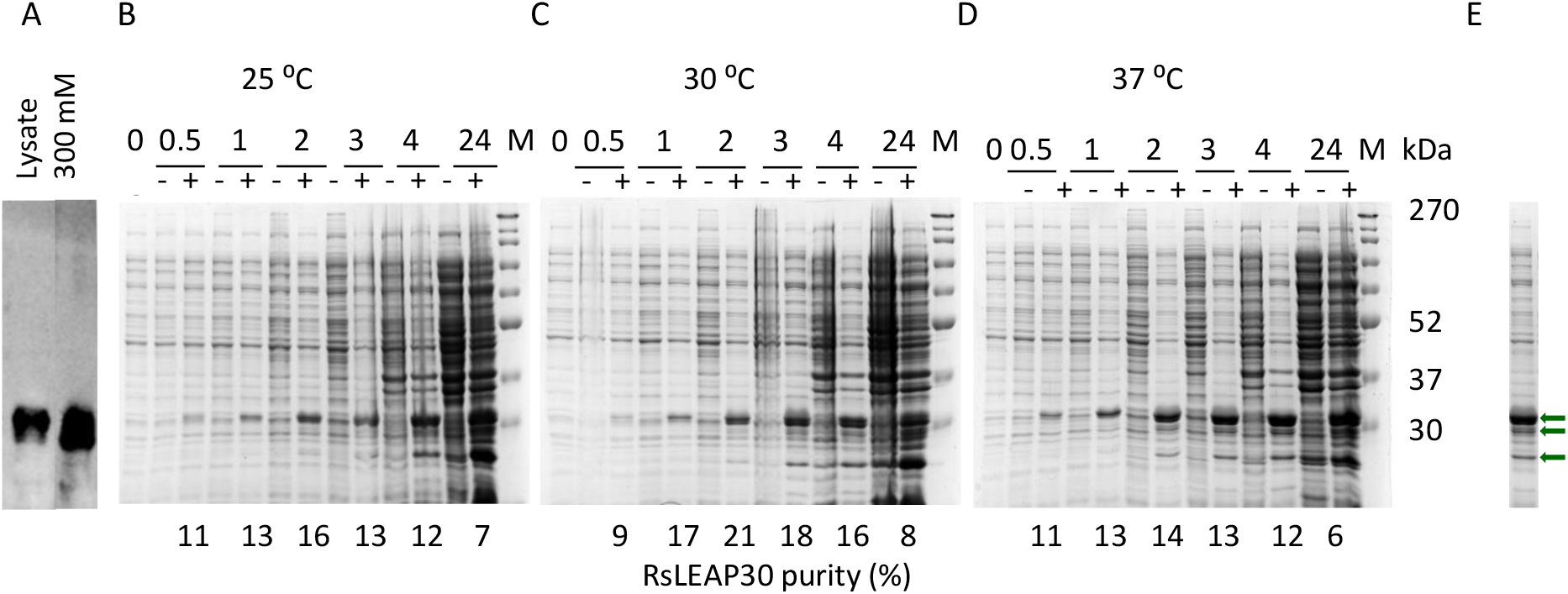
RsLEAP30 production in *E. coli* BL21 (DE3). **A)** Western blot of RsLEAP30 using anti-His antibody. Optimisation of RsLEAP30 production over a period of 24 hours at 25 °C **(B)**, 30 °C **(C)** and 37 °C **(D)**. Equal volumes of total protein extracts were loaded onto gels, which were subsequently stained and scanned according to a standardised procedure. The amounts of RsLEAP30 and total *E. coli* proteins were quantified using ImageJ. 0, samples before induction; “‒” conditions without IPTG addition; “+” induction with 1 mM IPTG; M, molecular markers (MWP06, NIPPON Genetics, Düren, Germany). The purity of RsLEAP30 is expressed as a ratio of RsLEAP30 to total *E. coli* proteins. **(E)** Three different RsLEAP30_H_, RsLEAP30_M_, and RsLEAP30_L_ isoforms are pointed by arrows.

As with GBP-RsLEAP30 production, the expression conditions (temperature and induction time) were optimised for RsLEAP30 production at a small scale. Unlike GBP-RsLEAP30, temperature was not a critical factor, as RsLEAP30 yield remained similar at 25 °C, 30 °C, and 37 °C (Figures 4B, 4C, 4D). The highest RsLEAP30 production was observed after 24 hours of induction (Supplementary Figure S3); however, this also resulted in a significant increase in *E. coli* host proteins at all temperatures. The highest RsLEAP30-to-host protein ratio was observed after 2 and 3 hours of induction at 30 °C (Figure 4C). The highest amount of RsLEAP30 was achieved at 37 °C after 3 and 4 hours (64 % and 76 %, respectively) with the similar RsLEAP30 / total *E. coli* proteins ratio (Supplementary Figure S3, Figure 4D).

Based on these results, the 3-hour induction at 37 °C was determined as the optimal condition for RsLEAP30 production in *E. coli* BL21(DE3).

### 3.6. Solubility test, purification and validation of RsLEAP30

To evaluate the solubility of RsLEAP30 under the optimised expression conditions, cell pellets collected at different time points were resuspended in Tris-HCl buffer, lysed, and separated into pellet and supernatant fractions. RsLEAP30 was consistently detected in the soluble fraction, regardless of incubation time, indicating its solubility in *E. coli* BL21 (DE3) (Figure 5).

**Figure 5.**
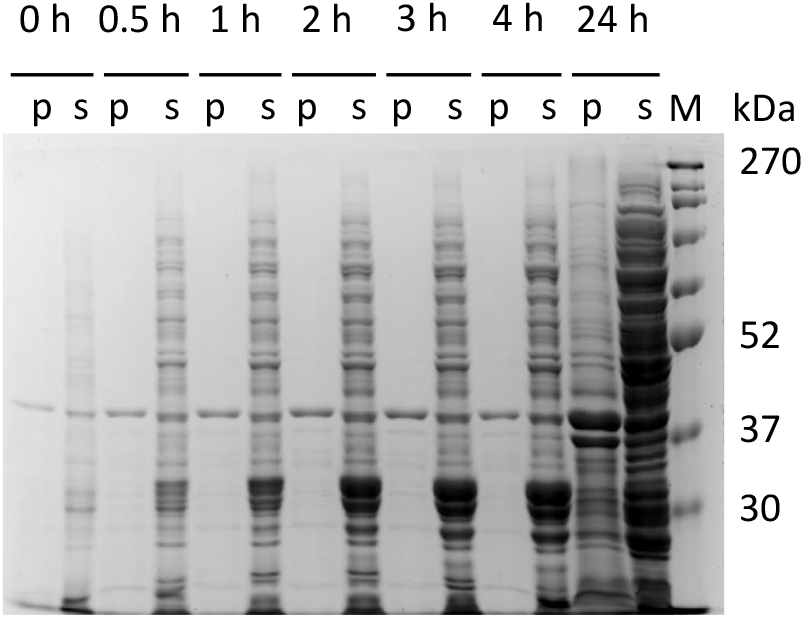
Solubility analysis of RsLEAP30 during a 24-hour incubation. For details, see Materials and Methods, section 2.5. M, molecular markers (MWP06, NIPPON Genetics, Düren, Germany); p, pellet; s, supernatant.

To refine the first purification step, IMAC elution conditions (10–300 mM imidazole) were optimised (Supplementary Figure S4). Stepwise elution with 30-50 mM imidazole resulted in the release of low molecular weight *E. coli* proteins, along with RsLEAP_L_ and RsLEAP_M_. At 75 mM imidazole, RsLEAP_M_ was eluted along with the initial release of RsLEAP_H_ isoform, which was more abundant. The highest yield of RsLEAP_H_ was obtained at imidazole concentrations above 200 mM (Supplementary Figure S4).

Based on these findings, an isocratic elution with 300 mM imidazole was chosen (Figure 6A), ensuring high RsLEAP30 recovery for all isoforms. A subsequent size-exclusion chromatography (SEC) step was performed to further separate RsLEAP_L_, RsLEAP_M_, and RsLEAP_H_ (Figure 6B). In contrast to GBP-RsLEAP30, RsLEAP30 was not detected in the IMAC FT fraction, indicating efficient column binding (Figures S4, 6B).

**Figure 6.**
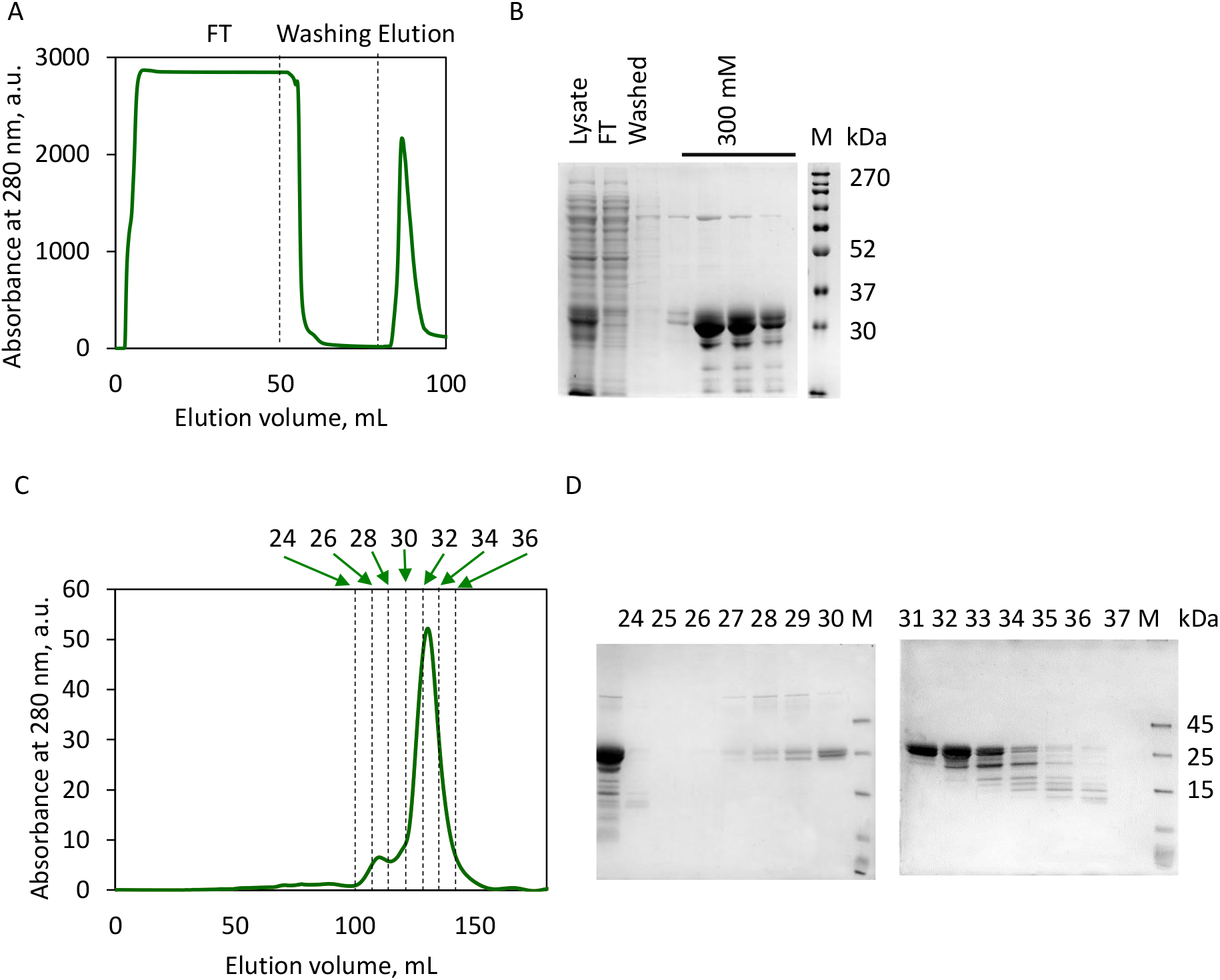
Purification of RsLEAP30. A) Representative IMAC chromatogram of RsLEAP30 elution. The peak containing RsLEAP30 was eluted with 300 mM imidazole. B) SDS-PAGE gel of the collected fractions from the IMAC. M, molecular markers (MWP06, NIPPON Genetics, Düren, Germany); FT, flow through. C) Size-exclusion chromatography (SEC) chromatogram of RsLEAP30. D) SDS-PAGE gel of the collected SEC fractions. M, molecular markers (MBS355494, San Diego, California).

Following SEC purification allowed separation of the fractions containing bands corresponding to the RsLEAP_L_, RsLEAP_M_ and RsLEAP_H_ isoforms (Figures 6C, 6D). The fraction from the centre of the elution peak (fraction 31) showed two closely migrating bands, corresponding to RsLEAP_H_ and RsLEAP_M_ on SDS-PAGE.

To determine their Mw precisely, SEC coupled with multi-angle light scattering (SEC-MALS) was performed. Analysis of fraction 31 revealed an average Mw of RsLEAP_H_ and RsLEAP_M_ of 24.64 ± 2.37 kDa, which deviates from the theoretical molecular weight of RsLEAP30 (28 kDa).

To further characterise these two protein isoforms, a bottom-up proteomic approach with mass spectrometry (MS) detection was employed. Sequence identification confirmed almost complete sequence coverage of RsLEAP_H_ (95 %) and RsLEAP_M_ (90 %). Since both isoforms exhibited a sequence coverage of more than 90, fraction 31 was considered suitable for further analyses.

Overall, from an initial culture volume of 1 L and a biomass of 5.6 g, a total yield of approximately 3.1 mg RsLEAP30 (from fraction 31) was obtained with a final purity reaching 98 % (Table 2).

**Table 2.** Purification table of RsLEAP30 from *E. coli* obtained from 1 L culture (from 5.6 g cell pellet).

| Fraction | Total protein content (mg) | RsLEAP30 purity (%)<sup>a</sup> | RsLEAP30 content (mg) | RsLEAP30 yield (%) |
| --- | --- | --- | --- | --- |
| Lysate | 142.2 | 50.0 | 71.2 | 100.0 |
| 300 mM imidazole<sup>b</sup> | 20.7 | 71.7 | 14.9 | 20.8 |
| Fraction 31 (SEC) | 3.20 | 98.0 | 3.14 | 4.4 |

<sup>a</sup> The purity of RsLEAP30 at each step of purification was estimated by SDS-PAGE.
<sup>b</sup> Polled 300 mM imidazole fraction shown in Figure 6B.

### 3.7. Secondary structure determination

The three-dimensional (3D) structure of RsLEAP30 is key to understanding its function. Therefore, we first constructed 3D models of RsLEAP30 under physiological conditions using AlphaFold2 software (Figure 7). Based on the three models obtained, RsLEAP30 shows a high propensity to adopt α-helix (Figure 7A), with a disorder content of 67 %, 39 %, and 43 % (Figure 7B), and low predicted alignment error (PAE) values (Supplementary Figure S5).

**Figure 7.**
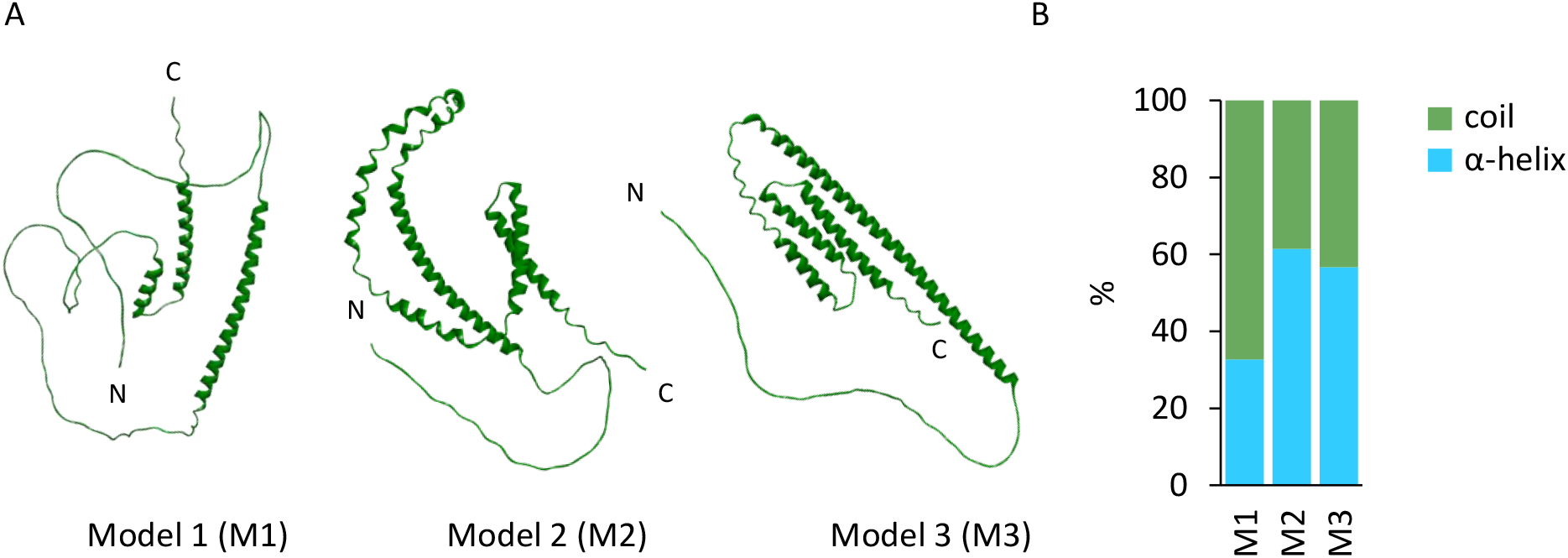
Three-dimensional models of RsLEAP30 according to AlphaFold2 (Jumper et al. 2021).

To investigate the presence of secondary structures in RsLEAP30 and the influence of pH on its structural conformation, we used circular dichroism (CD) spectroscopy in the far-ultraviolet (far-UV). CD spectra of RsLEAP30 were recorded under different pH conditions (pH 4–8) (Figure 8A). The spectra obtained at all five pH values tested showed an intense minimum at 201 nm, which is slightly shifted towards higher wavelengths, compared to the minimum at 195 nm, characteristic for random coils. Additionally, a slight minimum was observed at 222 nm, which is characteristic of α-helices.

**Figure 8.**
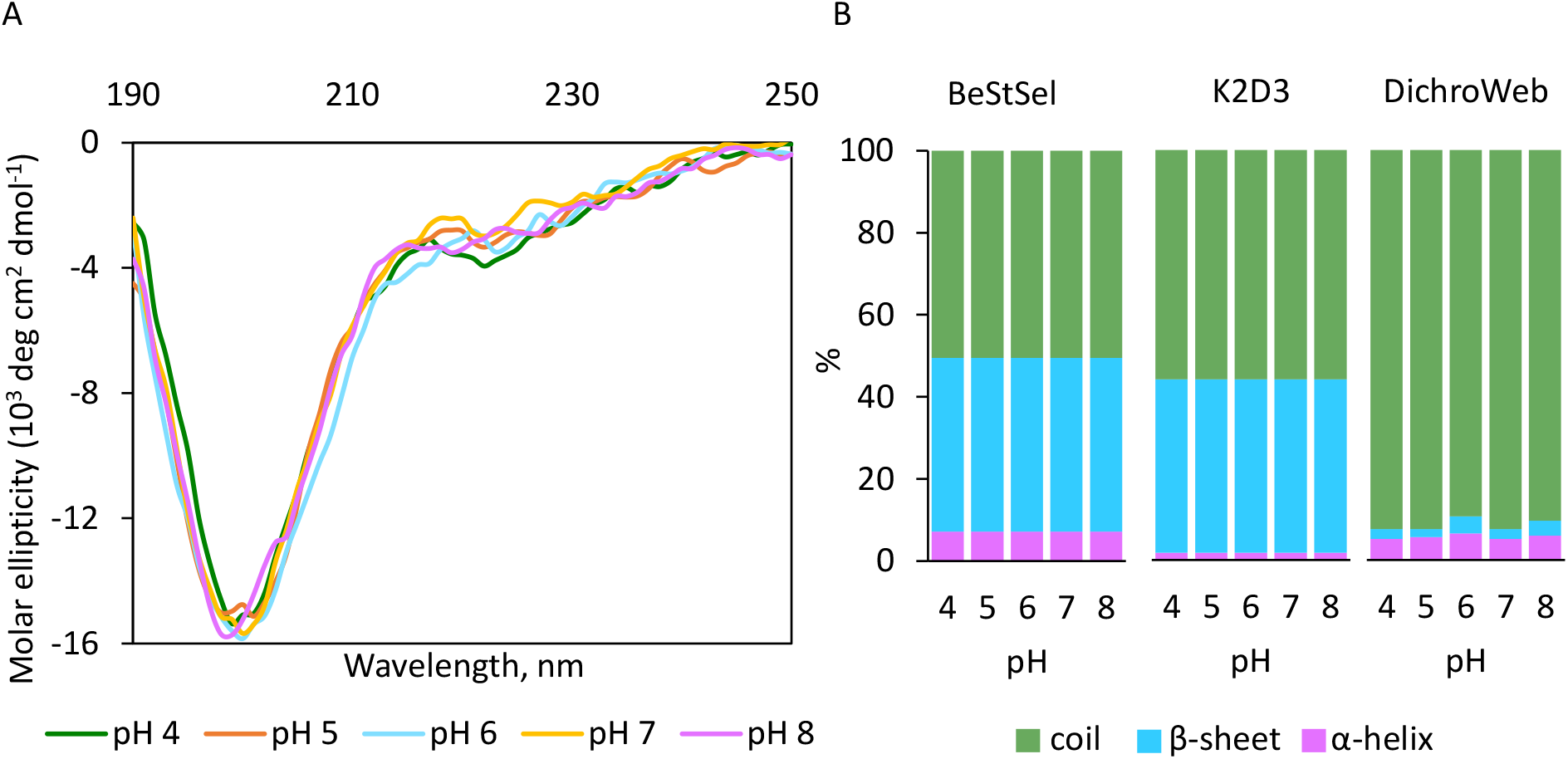
Effect of pH on the secondary structure of RsLEAP30 at 25°C, as monitored by CD spectroscopy. The percentage of secondary structure was evaluated using three software tools: BeStSel, DichroWeb, and K2D3.

To calculate the contribution of the different secondary structures, we employed three common software tools: BeStSel, DichroWeb, and K2D3 (Figure 8B). Each algorithm calculated different compositions of α-helices, β-sheets, and random coils. According to BeStSel, 7.2 %, 10 %, 41.6 %, and 41.3 % of the RsLEAP30 sequence was found in α-helical, β-turn, β-sheet, and disordered conformations, respectively. In contrast, DichroWeb determined that the majority of RsLEAP30 is disordered (∼ 94 % random coils), with only 4 % of the sequence folded into α-helices and ∼2% into β-sheets. The K2D3 software determined approximately 2 %, 43 %, and 55 % of RsLEAP30 adopting α-helical, β-sheet, and disordered structures, respectively (Figure 8B).

## Discussion

As a tertiary relict, *R. serbica* serves as an excellent model for studying vegetative desiccation tolerance, a phenomenon that is considered a pivotal step in the evolution of early land plants (Toldi et al. 2009). LEA proteins are a hallmark of vegetative desiccation tolerance. However, despite various proposed functions, their precise role remains unclear (Olvera-Carrillo et al. 2011; Dirk et al. 2020).

In this study, we report, for the first time, the recombinant expression of the LEA protein from the dicotyledonous, homoiochlorophyllous resurrection plant *R. serbica* using a bacterial expression system. Initially, we aimed to achieve high-yielding, tag-free RsLEAP expression in the *E. coli* BL21(DE3) system, given its simplicity, rapid growth, ease of genetic modification, and cost-effectiveness (Rosano and Ceccarelli 2014). However, this system presents certain challenges, including the formation of inclusion bodies and protein degradation (Bhatwa et al. 2021). Additionally, recombinant production of intrinsically disordered proteins (IDPs) is inherently difficult due to their structural plasticity and susceptibility to proteolysis and aggregation (Graether 2019). The presence of disordered and hydrophobic regions predisposes these proteins to the formation of inclusion bodies (Churion and Bondos 2012; Bhatwa et al. 2021).

To overcome these limitations, we integrated a fusion partner GBP in our initial approach, as it has been shown reduce the formation of inclusion body and increase the stability, solubility and yield of the target proteins (Rosano et al. 2019; Tepap et al. 2023). Consequently, we successfully obtained GBP-RsLEAP30 with an estimated molecular weight (Mw) of 46 kDa, achieving a yield of ∼15 % and a purity of ∼93 % (Figures 1–3; Table 1). The observed discrepancy in Mw, caused by retarded electrophoretic mobility compared to the computed Mw of GBP-RsLEAP30 (37 kDa), is more likely result of the high content of negatively charged amino acids (Glu 8%, Asp 8.3%), which reduces SDS binding efficiency, as previously reported (Kaufmann 1984; Weinreb 1996).

Despite numerous attempts to cleave the fusion partner with TEV protease under a variety of reaction conditions, we were unable to obtain native RsLEAP30. TEV protease is widely used for the cleavage of the fusion partners due to its high specificity, activity across diverse conditions and commercial availability. Since digestion of GBP-RsLEAP30 did not occur under optimal conditions for TEV proteolytic activity (despite successful cleavage of other TEV-site-containing recombinant proteins under the same conditions) we propose that GBP-RsLEAP30 adopts a conformation in which the recognition site (ENLYFQ/G) at the N-terminus is inaccessible, thereby preventing TEV binding. A similar explanation can be given for the weak interaction of GBP-RsLEAP30 with Ni-based resin during the first purification step using IMAC, as the His_8_ tag is also located in the N-terminal region, like GBP. This suggests that the N-terminus of GBP-RsLEAP30 may be structurally buried inside the protein.

Since TEV protease activity is not affected by certain detergents such as Triton X-100, Tween-20 and DDM (Sun et al. 2012), we attempted to induce conformational changes in GBP-RsLEAP30 to expose the TEV recognition site, but these efforts were unsuccessful. As TEV protease is a cysteine protease, we also tested different concentrations of reducing agents, including DTT, cysteine, GSH, and GSSG, but could not observe improvement in cleavage efficiency.

To circumvent the digestion issue, we designed a new construct that encodes RsLEAP30 with an His_6_ tag at the C-terminus, eliminating the need for a fusion partner. Given that RsLEAP30 is a highly polar protein, with 28 % polar and 34 % charged amino acids and a GRAVY index of ‒1.00 (Pantelić et al. 2022), the absence of a fusion partner is unlikely to affect its solubility in *E. coli*.

RsLEAP30 was expressed in *E. coli* BL21(DE3) cells at 37 °C for 3 hours as a soluble protein (Figure 4) and subsequently, it was purified using IMAC and SEC, as shown in Figure 6. RsLEAP30 exhibited less yield than GBP-RsLEAP30, due to the additional purification step (after Tables 1 and 2), while they achieved similar purities at the end, exceeding 93 %.

Recombinant DNA technology has already been successfully employed to produce LEA proteins in various organisms. For instance, six LEA proteins from the monocot poikilochlorophyllous resurrection species *Xerophyta schlechteri* were successfully expressed in *E. coli* (Artur et al. 2019). Similarly, three members of the LEA4 protein family were expressed in *E. coli* to generate specific antibodies for functional analyses in Arabidopsis mutants (Olvera-Carrillo et al. 2010). Eleven LEA proteins from *Dendrobium officinale* were also expressed in *E. coli* to investigate their roles in stress tolerance (Ling et al. 2016). In addition, LEA11 and LEA25 from Arabidopsis were produced in *E. coli* to study their potential interactions with membranes (Bremer et al. 2017). Recently, two recombinant LEA proteins from tardigrades and nematodes were expressed in *E. coli* and purified by the same two-step chromatographic methods (IMAC and SEC) employed in our study (Abe et al. 2024).

However, none of these reports provide quantitative data on yield and purity. To our knowledge, this is the first study to described the purity and yield of a recombinant LEA protein from resurrection plants. For comparison, in the case of other recombinant IDPs, such as human α-synuclein expressed in *E. coli*, the reported yield and purity were 30 % and 99 % respectively (Safari et al. 2020). Although the yield of RsLEAP30 was lower than expected for recombinant IDPs, purity was similar (98 %, Table 2). Our main objective was to obtain a highly purified protein for structural studies.

The experimentally determined molecular weights of GBP-RsLEAP30 and RsLEAP30, based on electrophoretic mobility, were higher than the theoretically expected values (Figures 1-6). This discrepancy can be attributed to the high proportion of negatively charged amino acids (see above), as well as potential post-translational modifications such as phosphorylation, which may influence electrophoretic migration (Hogema et al. 2002). Threonine, serine and tyrosine (amino acids that serve as phosphorylation sites) account for almost 18 % of RsLEAP30. Phosphorylation has already been shown to regulate certain LEA proteins (Alsheikh et al. 2003; Riera et al. 2004; Boudet et al. 2006). Indeed, variations in the degree of phosphorylation could explain the presence of three distinct mobility bands, corresponding to different RsLEAP30 isoforms (Figures 4-6). Additionally, negatively charged amino acids may mimic the phosphorylation-induced mobility shift on SDS-PAGE, which also contributes to slower migration (Lee et al. 2013).

Although mass spectrometry (MS) analysis of RsLEAP30_H_ confirmed a sequence identity of over 95 %, its Mw as determined by SEC-MALS, was slightly underestimated. Since SEC separates molecules based on hydrodynamic radius rather than molecular weight, this discrepancy can be attributed to a significant contribution of unstructured regions in RsLEAP30 (Figure 8). Proteins containing disordered regions often adopt an elongated conformation, leading to later elution and an underestimated Mw (Smith et al. 2024).

Our previous findings, based on five commonly used secondary structure prediction tools indicated that over 95% of the RsLEAP30 sequence has a high propensity to form α-helices (Pantelić et al. 2022). However, disorder predictors estimated that up to 85% of the sequence is intrinsically disordered. In this study, we performed 3D modelling using AlphaFold2, an advanced AI-based protein structure prediction tool that has significantly improved the accuracy of 3D protein structure predictions through deep learning techniques (Jumper et al. 2021). The resulting models indicated a disordered N-terminal region, while the segment spanning residues ∼60–243 was predominantly α-helical. According to HeliQuest web server predictions (Gautier et al. 2008), this region is expected to fold mainly into A-type α-helices, containing distinct positive, negative, and/or hydrophobic faces (Pantelić et al. 2022).

To experimentally characterise the secondary structure of RsLEAP30, we performed circular dichroism (CD) spectroscopy. RsLEAP30 was predicted to be localised in chloroplasts (Pantelić et al. 2022), where the stromal pH is ∼8, the luminal pH is 5–6, and the pH of the intermembrane space is 7.0–7.4. Since conformational changes in IDPs can be triggered by environmental factors such as pH (Uversky 2009), we examined the secondary structure of RsLEAP30 in a pH range of 4–8 to better understand the functional properties of these stress-associated proteins. Interestingly, CD spectroscopy revealed no significant changes in the secondary structure in this pH range, suggesting that RsLEAP30 is largely disordered (Figure 8).

Different CD analysis algorithms yielded different secondary structure predictions. BeStSel, which is optimised for β-rich and soluble proteins (Linhares et al. 2023), predicted a higher β-sheet content. In contrast, K2D3, which estimates α-helix and β-sheet content from its own theoretical reference dataset, reported increased α-helix content, which may be overestimated (Louis-Jeune et al. 2011). Additionally, when analysing our data with the DichroWeb server, we used dataset 7, which contains five spectra of denatured proteins that may or may not be disordered (Miles et al. 2022). Thus, the CD spectra analysis with DichroWeb (Figure 8) corroborate *in silico* predictions for the predominantly disordered nature of RsLEAP30 in aqueous solution. Similar results were reported for recombinantly produced LEA proteins from the monocot poikilochlorophyllous resurrection species *Xerophyta schlechteri* (Artur et al. 2019).

Based on *in silico* and spectroscopic data, we propose that the combination of disorder and α-helical structure in RsLEAP30 may be essential for its protective role in chloroplasts, particularly for maintaining membrane integrity under desiccation conditions. The α-helical regions of RsLEAP30 may interact electrostatically with the negatively charged phosphate groups of phospholipids via their positively charged side, while their hydrophobic side may interact with the fatty acid tails of membrane lipids. However, the inner envelope membrane of chloroplasts, etioplasts, and proplastids, as well as the thylakoid membrane, mainly consist neutral galactolipids and only 8–10 % of phospholipids (Candat et al. 2014). This composition suggests that interaction with membranes may be a minor role of RsLEAP30, while its interaction (via its A-type α-helices with alternating positive and negative sides) with desiccation-sensitive proteins within chloroplasts, particularly components of the photosynthetic electron transport chain, may be more important. This is particularly important since *R. serbica*, as a homoiochlorophyllous species, can rapidly resume photosynthesis after rehydration. Moreover, the presence of IDRs may allow RsLEAP30 to remain in a predominantly random coil conformation under normal physiological conditions, while adopting an α-helical structure during desiccation. Indeed, members of the LEA4 protein family in *Arabidopsis* have been shown to adopt an α-helical structure under water-limiting conditions, thereby preventing the inactivation and aggregation of lactate dehydrogenase *in vitro* (Cuevas-Velázquez et al. 2016). Furthermore, recombinant LEA proteins from the LEA1 family in *X. schlechteri* were found to adopt a fully α-helical conformation in hydrophobic acetonitrile solutions (Artur et al. 2019).

## Conclusion

Our study represents the first report on the recombinant production of a LEA protein from the dicot, homoiochlorophyllous resurrection plant *R. serbica*. Our results demonstrate that a combination of *in silico* and *in vitro* analyses can provide valuable insights into the structure of LEA proteins. We propose that the structure/function relationship of desiccation-induced LEA proteins in *R. serbica* plays a crucial role in its ability to rapidly restore photosynthetic components upon rehydration.

By elucidating the structure of RsLEAP30, we have provided a foundation for future research into its conformational plasticity and potential intracellular targets. Further investigation of RsLEAP30’s molecular interactions could enhance our understanding of the protective mechanisms underlying dehydration tolerance in resurrection plants. Such knowledge may be relevant for bioengineering strategies to improve drought resilience in crops.

## Acknowledgement

This research was funded by the Science Fund of the Republic of Serbia–RS (PROMIS project LEAPSyn-SCI, grant no. 6039663) and by the Ministry of Education, Science and Technological Development, the Republic of Serbia (Contract No. 451-03-136/2025-03/200042). A.P. wishes to acknowledge the support of COST Action CA21160 for approving Short-Term Scientific Missions in Ljubljana. We thank Dr Miloš Rokić for his suggestions regarding the RsLEAP30 purification.

