## Supplementary Figures for "Optimisation of *Ramonda serbica* LEA protein production in *Escherichia coli* and its secondary structure analysis": SFig1.pdf

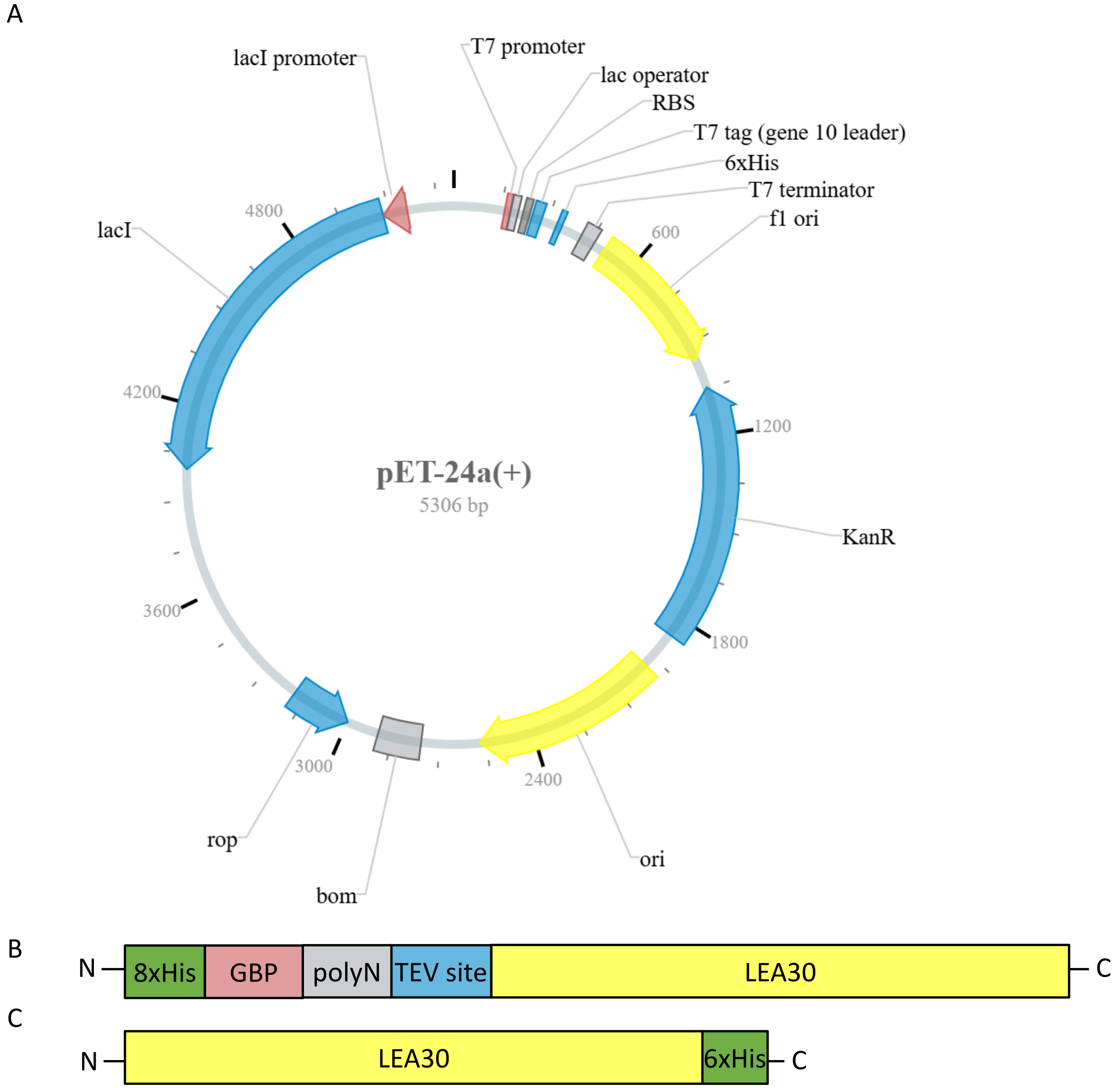


**Supplementary Figure S1.** Plasmid containing GBP-RsLEAP30 and RsLEAP30 constructs. **(A)** The pET-24a vector map, construct was inserted into XhoI and NdeI sites. **(B)** Schematic illustration of GBP-RsLEAP30 construct. **(C)** Schematic illustration of RsLEAP30 construct.
