## Supplementary Figures for "Optimisation of *Ramonda serbica* LEA protein production in *Escherichia coli* and its secondary structure analysis": SFig2.pdf

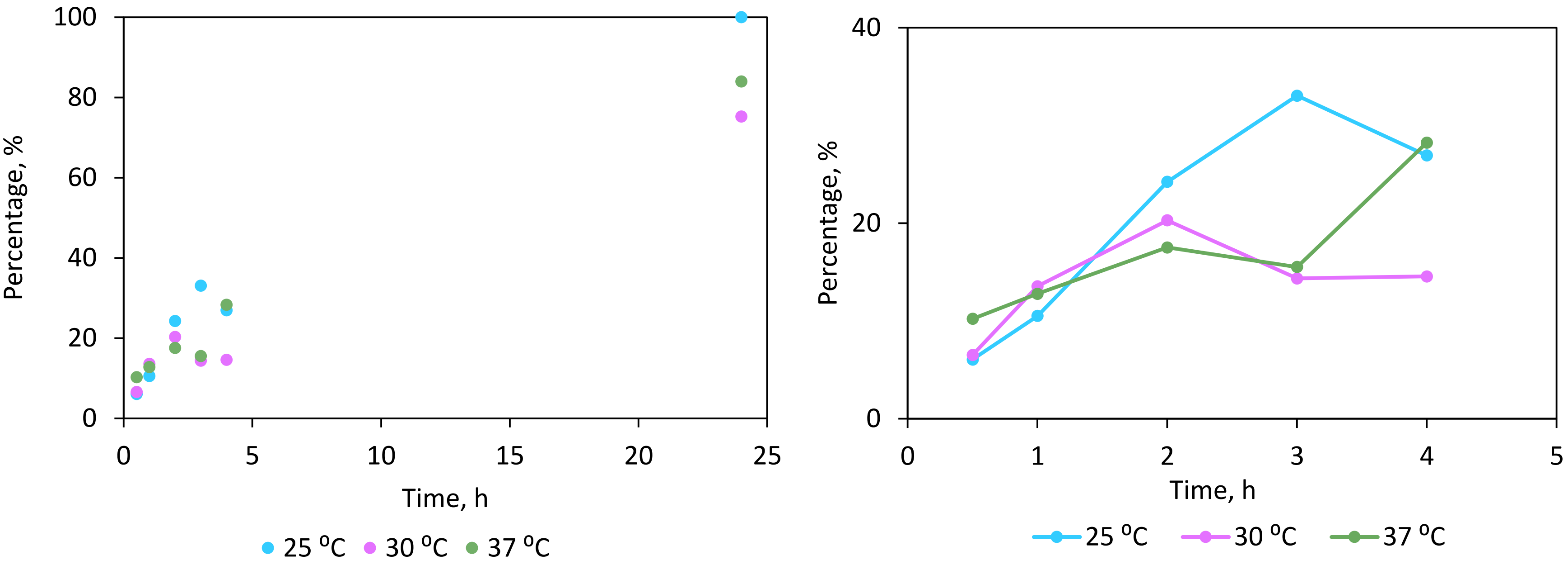


**Supplementary Figure S2.** The production rate of GBP-RsLEAP30, during 24 h, at three temperatures (25 °C, 30 °C and 37 °C). The amount of the produced GBP-RsLEAP30 is expressed as a percentage, where 100 % represents the highest amount of GBP-RsLEAP30 achieved at 25 °C after 24 h.
