## Supplementary Figures for "Optimisation of *Ramonda serbica* LEA protein production in *Escherichia coli* and its secondary structure analysis": SFig4.pdf

A

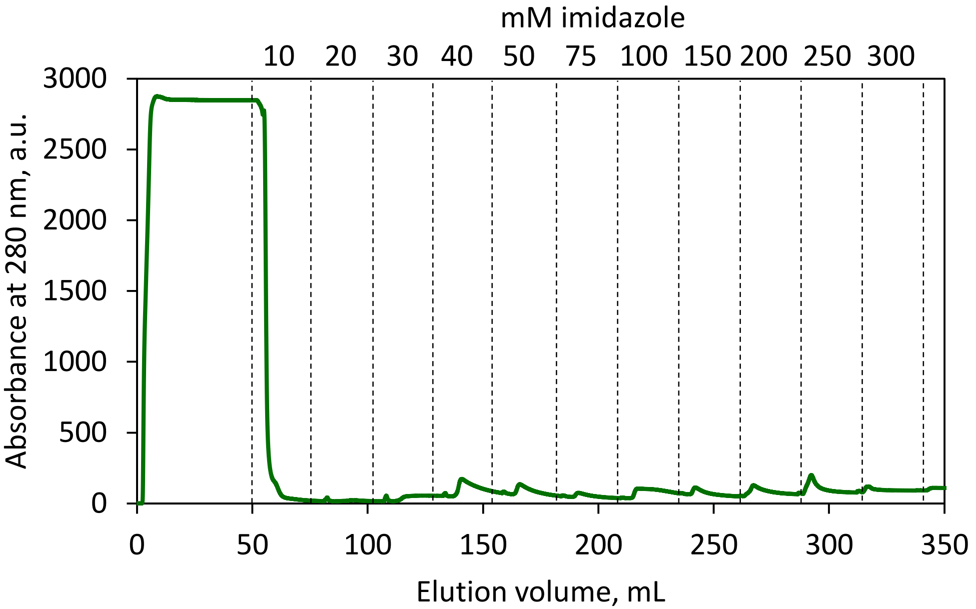


B

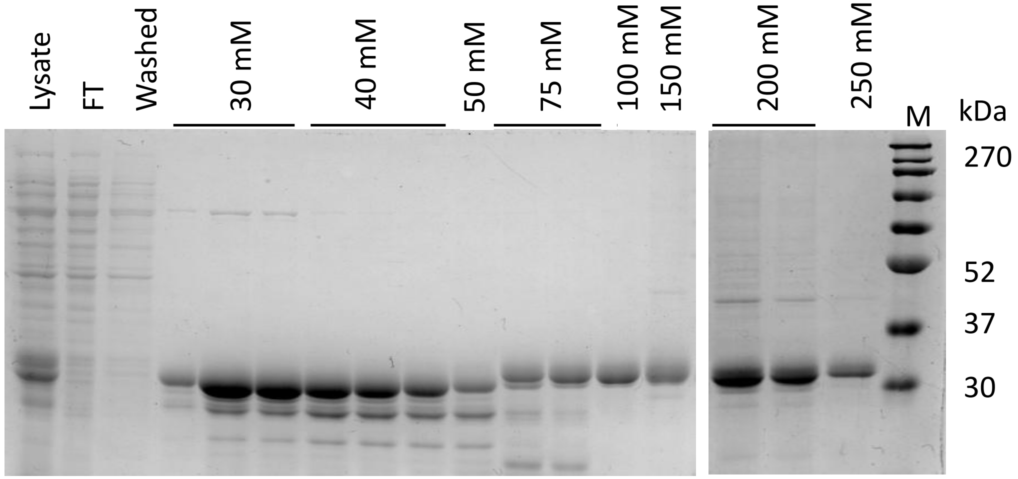


**Supplementary Figure S4.** IMAC optimisation of RsLEAP30 **(A)** and SDS-PAGE gel of the collected fractions **(B)**. M, molecular markers (BlueEasy Prestained Protein Marker, MWP06, NIPPON Genetics, Düren, Germany). 10-300 mM represent the imidazole concentration in the elution buffer.
