## Supplementary Figures for "Optimisation of *Ramonda serbica* LEA protein production in *Escherichia coli* and its secondary structure analysis": SFig5.pdf

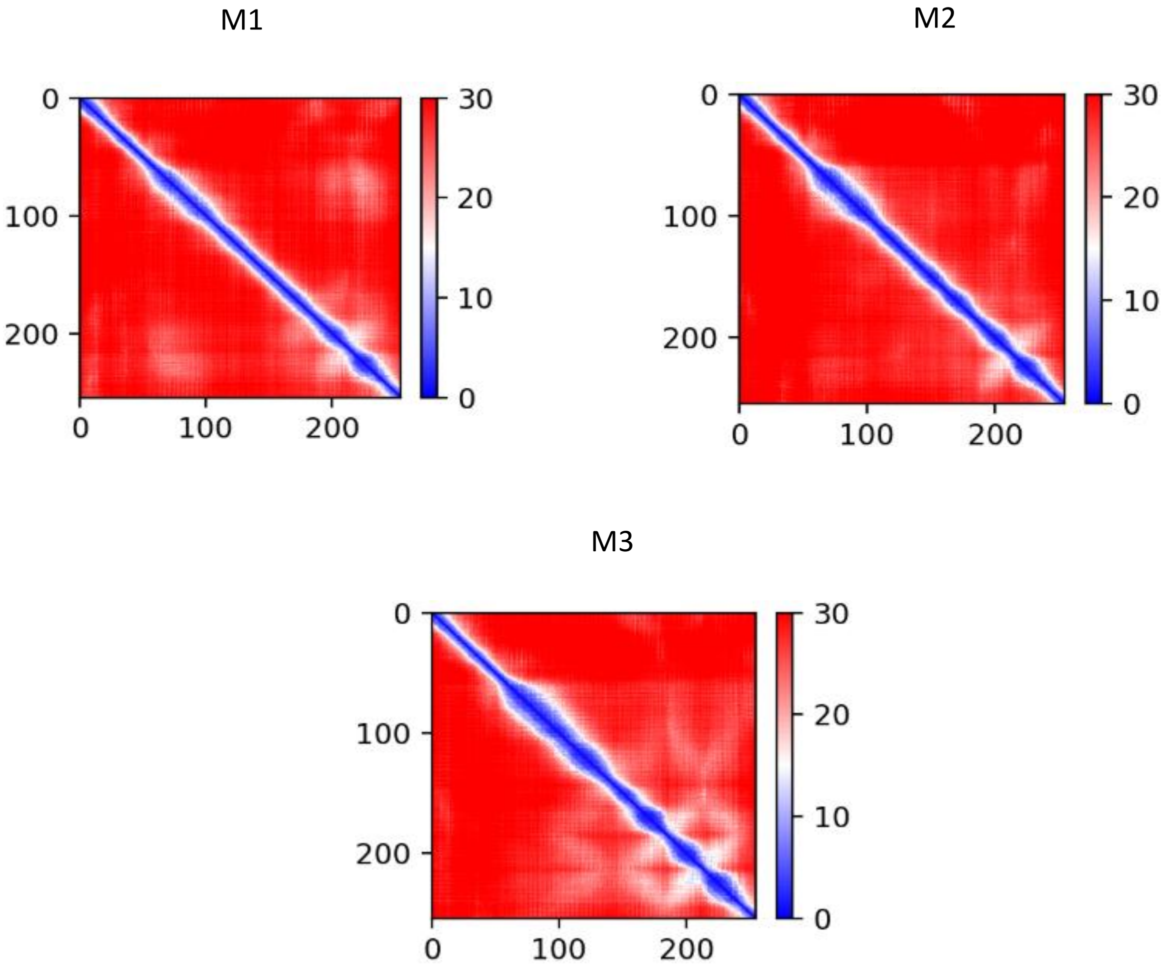


**Supplementary Figure s5.** The Predicted Aligned Error (PAE) of three AlphaFold structure of RsLEAP30., shows the predicted relative position error for each residue in the sequence, with low-confidence values in red.
